# SMARTer single cell total RNA sequencing

**DOI:** 10.1101/430090

**Authors:** Verboom Karen, Everaert Celine, Bolduc Nathalie, Livak J. Kenneth, Yigit Nurten, Rombaut Dries, Anckaert Jasper, Venø T Morten, Kjems Jørgen, Speleman Frank, Mestdagh Pieter, Vandesompele Jo

## Abstract

Single cell RNA sequencing methods have been increasingly used to understand cellular heterogeneity. Nevertheless, most of these methods suffer from one or more limitations, such as focusing only on polyadenylated RNA, sequencing of only the 3’ end of the transcript, an exuberant fraction of reads mapping to ribosomal RNA, and the unstranded nature of the sequencing data. Here, we developed a novel single cell strand-specific total RNA library preparation method addressing all the aforementioned shortcomings. Our method was validated on a microfluidics system using three different cancer cell lines undergoing a chemical or genetic perturbation. We demonstrate that our total RNA-seq method detects an equal or higher number of genes compared to classic polyA[+] RNA-seq, including novel and non-polyadenylated genes. The obtained RNA expression patterns also recapitulate the expected biological signal. Inherent to total RNA-seq, our method is also able to detect circular RNAs. Taken together, SMARTer single cell total RNA sequencing is very well suited for any single cell sequencing experiment in which transcript level information is needed beyond polyadenylated genes.

## Introduction

To understand the complexity of life, knowledge of cells as fundamental units is key. Recently, technological advances have emerged to enable single cell RNA sequencing (RNA-seq). To date, droplet and split-pool ligation based methods capture thousands of single cells, providing new insights in cellular heterogeneity and rare cell types (1–5). The main drawback of these methods is that analyses are typically confined to gene expression of only (3’ ends of) polyadenylated transcripts. More complex analyses with respect to alternative splicing, allele specific expression, mutation analysis, assembly of (novel) transcripts, circular RNA (circRNA) quantification and post-transcriptional regulation, require full-length and full-transcriptome methods. Moreover, sequencing a large number of cells is often compromising sequencing depth, resulting in low coverage per cell and detection of only the most abundant transcripts (6). In contrast to these droplet based methods, microfluidic chip and flowcytometry based platforms typically capture fewer cells, but are able to sequence entire transcripts and detect a substantially higher number of genes per cell providing a more complete view of the complexity and richness of single cells’ transcriptomes (7, 8). Of note, most single cell RNA-seq studies assess only 3’ end polyadenylated (polyA[+]) transcripts, ignoring non-polyadenylated (polyA[-]) transcripts. Since a substantial part of the human transcriptome is non-polyadenylated, various RNA types including circRNAs, enhancer RNAs, histone RNAs, and a sizable fraction of long non-coding RNAs (lncRNAs) are not quantified using these classic methods (9–11). In order to study polyA[-] transcripts at the single cell level, total RNA-seq workflows were developed (12–14). While in principle both polyA[+] and polyA[-] transcripts are converted into a sequencing-ready library using random primer mediated reverse transcription, these methods suffer from one or more of the following limitations: the strand-orientation information is lost and a high percentage of reads map to ribosomal RNA (rRNA). Therefore, new methods circumventing these limitations are warranted. A rRNA depletion step is essential as up to 95 % of the total RNA content in a mammalian cell consists of rRNA. Moreover, to discriminate sense and antisense overlapping transcripts, stranded sequencing is crucial; at least 38 % of the annotated transcripts in cancer cells have antisense expression (15). Here, we developed an easy to use and efficient single cell total RNA-seq workflow based on the SMARTer Stranded Total RNA-Seq Kit – Pico Input Mammalian including a ribodepletion step at the cDNA level. We ported the method to Fluidigm’s C1 single cell microfluidics instrument, but other systems should be equally feasible (e.g. FACS). In total, 391 cells from 3 different human cancer cell lines in 3 experiments were sequenced with a total sequencing depth of 1487 million reads. Using our novel method, we consistently observe less than 3 % of ribosomal reads and we detect more than 5360 genes by at least four reads, including novel genes, polyA[-] genes and circRNAs.

## Material and Methods

### Cell lines

The neuroblastoma cell line NGP is a kind gift of prof. R. Versteeg (Amsterdam, the Netherlands). Cells were maintained in RPMI-1640 medium (Life Technologies, 52400–025) supplemented with 10 % fetal bovine serum (PAN Biotech, P30–3306), 1 % of L-glutamine (Life Technologies, 15140–148) and 1 % penicillin/streptomycin (Life Technologies, 15160–047) (referred to as complete medium) at 37 °C in a 5 % CO_2_ atmosphere. Short tandem repeat genotyping was used to validate cell line authenticity prior to performing the described experiments and mycoplasma testing was done on a monthly basis.

### Cell cycle synchronization and nutlin-3 treatment of NGP cells

NGP cells were synchronized using serum starvation prior to nutlin-3 treatment. First, cells were seeded at low density for 48 hours in complete medium. Then, cells were refreshed with serum-free medium for 24 hours. Finally, the cells were treated with either 8 μM of nutlin-3 (Cayman Chemicals, 10004372, dissolved in ethanol) or vehicle. Cells were trypsinized (Gibco, 25300054) 24 hours post treatment and harvested for single cell analysis, bulk RNA isolation and cell cycle analysis.

### Cell cycle analysis

Four million cells were washed with PBS (Gibco, 14190094) and the pellet was resuspended in 300 µl PBS. Next, 700 µl of 70 % ice-cold ethanol was added dropwise while vortexing to fix the cells. Cells were stored at -20 °C for at least 1 hour. After incubation, cells were washed with PBS and the pellet was resuspended in 1 ml PBS containing RNAse A (Qiagen, 19101) at a final concentration of 0.2 mg/ml. After 1 hour incubation at 37 °C, propidium iodide (BD biosciences, 556463) was added to a final concentration of 40 µg/ml. Samples were loaded on a S3 cell sorter (Bio-Rad) and analyzed using the FlowJo v.10 software.

### RNA isolation and cDNA synthesis

Total RNA was isolated using the miRNeasy mini kit (Qiagen, 217084) with DNA digestion on-column according to the manufacturer’s instructions. RNA concentration was measured using spectrophotometry (Nanodrop 1000, Thermo Fisher Scientific). cDNA was synthesized using the iScript Advanced cDNA synthesis kit (Bio-Rad, 1708897) using 500 ng RNA as input in a 20 µl reaction. cDNA was diluted to 2.5 ng/µl with nuclease-free water prior to RT-qPCR measurements.

### Reverse transcription quantitative PCR

PCR mixes containing 2.5 µl 2x SsoAdvansed SYBR qPCR supermix (Bio-Rad, 04887352001), 0.25 µl each forward and reverse primer (5 µM, IDT), and 2 µl diluted cDNA (5 ng total RNA equivalents) were analyzed on the LightCycler480 instrument (Roche) using two replicates. Expression levels were normalized using expression data of four stable reference genes (SDHA, YWHAZ, TBP, HPRT1). RTqPCR data was analyzed using the qbase+ software v3.0 (Biogazelle). Primer sequences are available in Supplementary Table 1.

### Single cell total RNA library preparation of nutlin-3 treated NGP cells

Cells were washed with PBS and pellets of vehicle treated cells were resuspended and incubated in 1 ml pre-warmed (37 °C) cell tracker (CellTracker Green BODIPY Dye, Thermo fisher Scientific, C2102) for 20 minutes at room temperature. After incubation, cells were washed in PBS and resuspended in 1 ml wash buffer (Fluidigm, 100–6201). An equal number of stained (vehicle treated) and non-stained (nutlin-3 treated) cells were mixed and diluted to 300,000 cells per ml. Suspension buffer (Fluidigm) was added to the cells in a 3:2 ratio and 6 µl of this mix of was loaded on a primed C1 Single-Cell Open App IFC (Fluidigm, 100–8134) designed for medium-sized cells (10–17 µm). Cells were captured using the ‘SMARTer single cell total RNA-seq’ script deposited in Script Hub (Fluidigm). Upon capture, cells were visualized using the Axio Observer Z1 (Zeiss). Sequencing libraries were generated using the C1 running the ‘SMARTer single cell total RNA-seq’ script deposited on Script Hub. In short, the SMARTer Stranded Total RNA-Seq Kit v2 – Pico Input Mammalian (Pico v2, total RNA, Takara, 634413) was used to synthesize cDNA with following modifications. Cells were fragmented and lysed by loading 7 µl of 10x reaction mix [2.3 µl SMART Pico Oligo Mix v2, 6 µl 5x first-strand buffer, 1 µl 20x C1 loading reagent (Fluidigm), 3 µl lysis mix (19 µl 10x lysis buffer, 1 µl RNAse inhibitor (40 U/µl)), 1 µl 1/1000 diluted ERCC spikes (Ambion, 4456740), 6.7 µl water] and incubating the cells at 85 °C for 6 minutes (to lyse cells and fragment RNA) followed by 2 minutes at 10 °C. Next, 8 µl first strand master mix [1 µl C1 loading reagent, 4 µl 5x first-strand buffer, 0.9 µl RNAse inhibitor (40 U/µl), 3.5 µl SMARTScribe reverse transcriptase (100 U/µl), 7.9 µl SMART TSO Mix v2 (from Takara kit, 634413), 2.7 µl water] was loaded and incubated at 42 °C for 90 minutes followed by 70 °C for 10 minutes. Finally, a PCR master mix for each well was made [1 µl 20x loading reagent, 2 µl 2.4 µM forward primer (Takara, 634413), 2 µl 2.4 µM reverse primer, 13.1 µl 1.5x PCR mix (1050 µl 2x SeqAmp CB buffer, 42 µl SeqAmp DNA polymerase, 308 µl water)] and 5 µl of each of these mixes was loaded in the harvest wells of the IFC. The samples were incubated for 1 minute at 94 °C followed by 11 PCR cycles (30 s at 98 °C, 15 s at 55 °C, 30 s at 68 °C) and 2 minutes at 68 °C. Following this initial cDNA amplification, 12 wells were pooled per tube using 8 µl of cDNA per cell. Next steps of the library prep were performed according to manufacturer’s instructions with minor modifications. 13 PCR cycles were used for PCR2 and a 1:1 ratio was used for beads cleanup after PCR2. Next, the samples were resuspended in 22 µl 5 mM tris buffer (from kit) and 20 µl was used to perform a second beads cleanup using a 0.9:1 ratio. Finally, the samples were resuspended in 12 µl tris buffer and the quality was determined on the Fragment Analyzer (Advanced Analytical). Of note, the protocol can be modified using the single cell specific version of the kit, released by Takara (SMART-Seq Stranded Kit, 634442) after we had completed our experiment.

### Single cell polyA[+] RNA library preparation of nutlin-3 treated NGP cells

Vehicle treated cells were stained with cell tracker as described above. An equal number of stained (vehicle treated) and non-stained (nutlin-3 treated) cells were mixed and diluted to 300,000 cells per ml. Suspension buffer was added to the cells in a 3:2 ratio and 6 µl of this mix of was loaded on a primed C1 Single-Cell Auto Prep Array for mRNA Seq (Fluidigm, 100–6041) designed for medium-sized cells (10–17 µm). Single cell polyA[+] RNA sequencing on the C1 was performed using the SMART-Seq v4 Ultra Low Input RNA Kit for the Fluidigm C1 System (SMART-Seq v4, polyA[+] RNA, Takara, 635026) according to manufacturer’s instructions. One microliter of the ERCC spike-in mix was diluted in 999 µl loading buffer to get a 1/1000 dilution of the ERCC spikes. One microliter of this dilution was added to the 20 µl lysis mix. The quality of the cDNA was checked for 11 random single cells on the Fragment Analyzer. The concentration of the cells was measured using the quantifluor dsDNA kit (Promega, E2670) and glomax (Promega) according to manufacturer’s instructions. The samples were 1/5 diluted in C1 harvest reagent (Fluidigm). Next, library prep was performed using the Nextera XT library prep kit (Illumina, FC-131–1096) according to manufacturer’s instructions, followed by quality control on the Fragment Analyzer.

### Library sequencing

The polyA[+] and total RNA libraries were quantified using the KAPA library quantification kit (Roche) and libraries were diluted to 4 nM. The polyA[+] RNA library and total RNA library were pooled in a 1/4 ratio and 1.5 pM of the pooled library was single-end sequenced on a NextSeq 500 (Illumina) with a read length of 75 bp and a total sequencing read depth of 274 million reads, combining single cell polyA[+] and total RNA libraries to prevent inter-run bias. A median sequencing read depth of 0.81 and 3.67 million reads per cell was reached for the single cell polyA[+] and total RNA libraries, respectively. In addition, 1.3 pM of the total RNA library was also sequenced in 2×75 paired-end sequencing run mode on the NextSeq 500, yielding 327 million reads and a median sequencing read depth of and 4.04 million per cell. The fastq data is deposited in GEO (GSE119984).

### Sequencing data quality control

While single-end sequencing libraries do not require pre-trimming, the paired-end libraries were trimmed using cutadapt (v.1.16) (16) to remove 3 nucleotides of the 5’ end of read 2. To assess the quality of the data, the reads were mapped using STAR (v.2.5.3) (17) on the hg38 genome including the full ribosomal DNA (45S, 5.8S and 5S) and mitochondrial DNA sequences. The parameters of

STAR were set to retain only primary mapping reads. Using SAMtools (v1.6) (18), reads mapping to the different nuclear chromosomes, mitochondrial DNA and rRNA were extracted and annotated as exonic, intronic or intergenic. In contrast to the unstranded nature of polyA[+] SMART-seq data, the total RNA SMARTer-seq data is stranded and processed accordingly. Gene body coverage was calculated using the full Ensembl (v91) (19) transcriptome. The coverage per percentile was calculated, followed by a loess regression fit.

### Quantification of Ensembl and LNCipedia genes

Genes were quantified by Kallisto (v.0.43.1) (20) using both Ensembl (v.91) (19) extended with the ERCC spike sequences and LNCipedia (v.5.0) (21). The strandedness of the total RNA-seq reads was considered by running the –rf-stranded mode. Subsampling 1 million reads (polyA[+] RNA libraries) or 1, 4 or 8 million reads (total RNA libraries) was performed by seqTK (v.1.2) followed by Kallisto quantification. Further processing was done with R (v.3.5.1) making use of tidyverse (v.1.2.1). To measure the biological signal we first performed differential expression analysis between the treatment groups using DESeq2 (v.1.20.0) (22) in combination with Zinger (v.0.1.0) (23). To identify enriched gene sets a fsgea (v.1.6.0) analysis was performed, calculating enrichment for the hallmark gene sets retrieved from MSigDB (v.6.2).

### Circular RNA detection

CircRNAs were detected using the deeper sequenced paired-end sequencing data.Trim_galore (v.0.4.1) was used to trim adaptor sequences, perform quality filtering and remove 3 nucleotides from the 5’ end of read 2. Subsequently, reads from all samples were combined, adding originating sample name to read names for later splitting of data. The combined data was used for circRNA detection using find_circ (v.1) (24) using the reads2sample (find_circ.py –r) option to allow circRNA detection on the combined dataset while dividing out the contribution from each sample in the output. Only circRNAs with unique mapping on both anchors were accepted. Human genome hg19 was used for circRNA analysis. CircRNAs were annotated with host gene names from RefSeq (release 75) and circBase IDs from circbase.org. The Database for Annotation, Visualisation and Integrated Discovery (DAVID, v.6.8) (25, 26) was used for Gene Ontology (GO) analysis for the circRNA host genes using biological processes (BP) and molecular function (MF). P-value < 0.05 was used for statistical significance.

### Single cell transcriptome assembly

A transcriptome per cell was created by combining STAR (v.2.5.3) and Stringtie (v.1.3.0) (27), using the deeper sequenced paired-end sequencing data. The parameters of Stringtie were set to require a coverage of 1. These single cell transcriptomes were merged with the Ensembl (v.91) transcriptome as a reference. From the merged multi-cell transcriptome, only multi-exonic genes with a minimum length of 200 nt were retained. To define the set of novel genes, genes annotated in Ensembl (19) or LNCipedia (v.5.0) (21) were filtered out. All genes in this novel multi-cell transcriptome were quantified using Kallisto on single-end subsampled data (1, 4 or 8 million reads per cell). Genes with an estimated count higher than 1 were retained.

## Results

### Principle of SMARTer single cell total RNA sequencing

We developed a single cell total RNA-seq protocol for unbiased, full transcript and strand-specific analysis of both polyadenylated and non-polyadenylated transcripts from mammalian cells. The method is based on the SMARTer Stranded Total RNA-Seq Kit v2 – Pico Input Mammalian (Pico v2, total RNA), a Takara kit that is meant for low input bulk total RNA-seq. The library preparation method employs random primers and a template switching mechanism to capture full transcript fragments of both polyadenylated (polyA[+]) and non-polyadenylated (polyA[-]) transcripts. Unwanted ribosomal cDNA is removed using Pico v2 probes, specific to mammalian rRNA. After successfully porting the bulk library prep protocol to Fluidigm’s C1 single cell instrument, we assessed the performance of the single cell total RNA-seq protocol through three distinct experiments in which nutlin-3, JQ1 or doxycycline was used to treat NGP, SK-N-BE-2C, and SHSY5YMYCN-TR neuroblastoma cell lines, respectively (with vehicle treated cells as control) (Figure 1). In addition, we performed matched single cell polyA[+] RNA-seq as a reference using cells from the same pool. While all experiments were successful, we focus our analyses and performance assessment on the NGP data. In this experiment, the treated and control cells were processed in the same microfluidic chip (preventing possible chip bias), the highest number of cells were captured, and the highest sequencing depth was reached.

**Figure 1.**
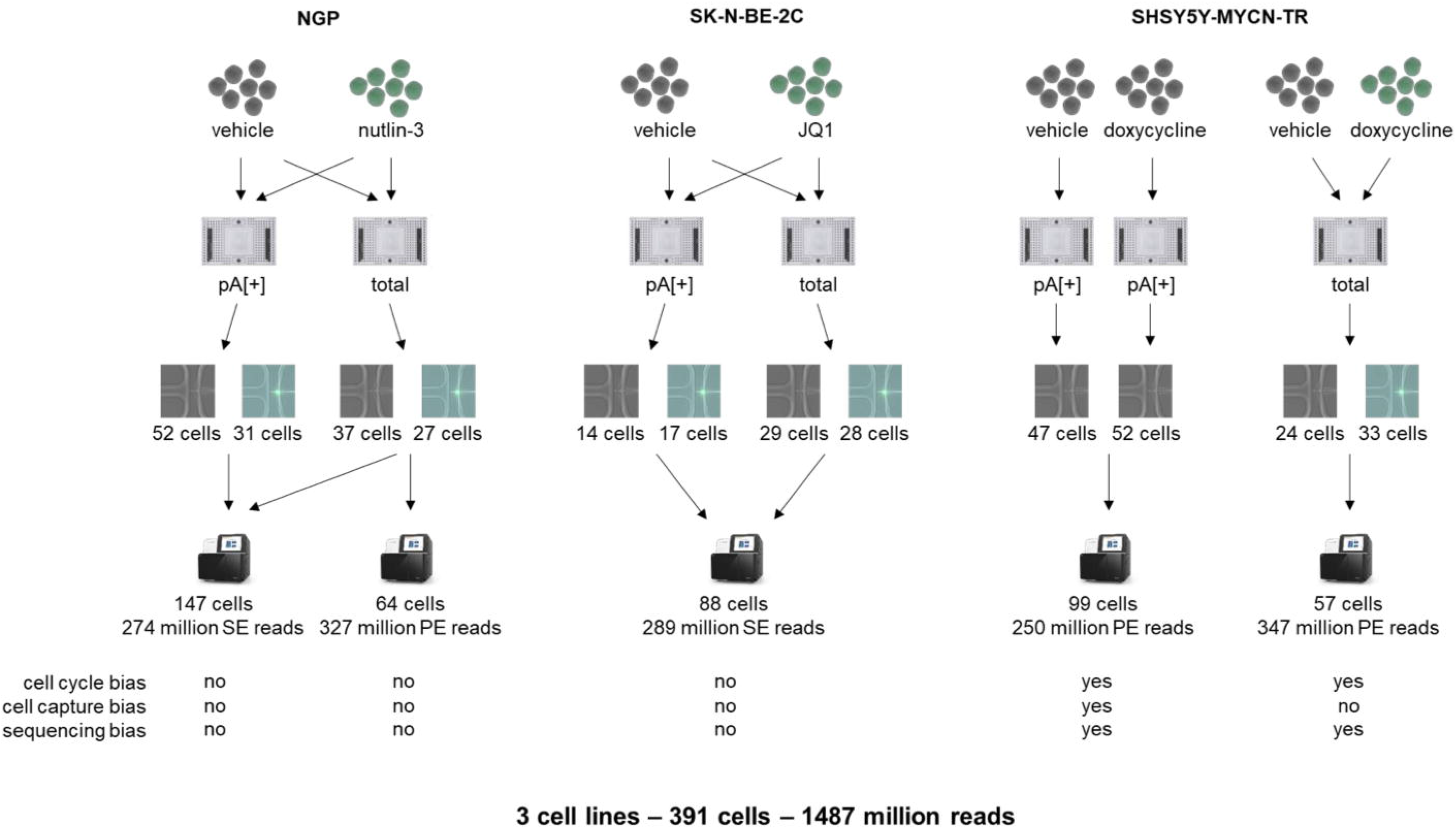
Overview of experimental set-up.

### SMARTer single cell total RNA sequencing yields high-quality data

In single cell sequencing experiments, it is important to prevent or limit potential biases that mask true biological differences. In particular, the cell cycle state is a known confounder (28). Therefore, we synchronized the cells through serum starvation for 24 hours. Upon synchronization, 80.3 % of the NGP cells showed an arrest at the G0/G1 stage compared to only 53.3 % for non-synchronized NGP cells (Supplementary Figure 1 A-B). Subsequently, the synchronized NGP cells were treated for 24 hours with vehicle or nutlin-3, the latter known to release TP53 from its negative regulator MDM2. As expected, nutlin-3 treatment resulted in cell cycle arrest (Supplementary Figure 1 C-D). To prevent possible C1 batch effects (29), vehicle treated NGP cells were stained and loaded together with the non-stained nutlin-3 treated cells on the same C1 chip. Based on the fluorescent label and the transparency of the C1 system, vehicle and nutlin-3 treated cells were discriminated by fluorescence microscopy. By loading two C1 chips, one for polyA[+] RNA and one for total RNA library preparation, we captured 31 and 27 nutlin-3 treated versus 52 and 37 vehicle treated single cells, respectively. High-quality cDNA libraries of polyA[+] and total RNA were generated using the SMART-Seq v4 Ultra Low Input RNA Kit for the Fluidigm C1 System (SMART-Seq v4, polyA[+]) and our novel SMARTer single cell total RNA-seq protocol, respectively (Supplementary Figure 1 E-F). ERCC spike-in molecules were added for external quality control in the lysis mix (Supplementary Figure 2). For the recovered spikes (with a concentration in the original mix of at least 10 attomoles/µl), linear models were calculated (Supplementary Figure 3), retrieving similar R^2^ values for the polyA[+] RNA and total RNA library preparation protocol (Supplementary Figure 4). The transcripts detected in the polyA[+] libraries were somewhat shorter compared to the total RNA libraries (Supplementary Figure 5). In addition, the total RNA-seq libraries show a more uniform transcript coverage (Supplementary Figure 6).

As expected, a higher fraction of reads mapped to nuclear rRNA in the total RNA-seq libraries compared to the polyA[+] RNA libraries (average of 2.739 % [2.488, 2.990; 95 % confidence interval (CI)] vs. 0.031 % [0.026, 0.035; 95 % CI], respectively). Nevertheless, the fraction of nuclear rRNA is very low in the total RNA libraries considering the use of random priming data (Figure 2A). Furthermore, the single cell total RNA libraries contain more intronic (27.99 % [25.06, 30.91; 95 % CI] vs. 11.87 % [10.14, 13.60; 95 % CI]) and intergenic (5.38 % [5.00, 5.76; 95 % CI] vs. 2.90 % [2.54, 3.26; 95 % CI]) nuclear reads compared to polyA[+] RNA libraries (Figure 2B). Non-polyadenylated histone genes are highly abundant in the total RNA libraries, while low or absent in the polyA[+] libraries, confirming the validity of our single cell total RNA-seq workflow (Supplementary Figure 7). Equal results were obtained for the SK-N-BE-2C, and SHSY5Y-MYCN-TR cell lines (Supplementary Figure 8).

**Figure 2.**
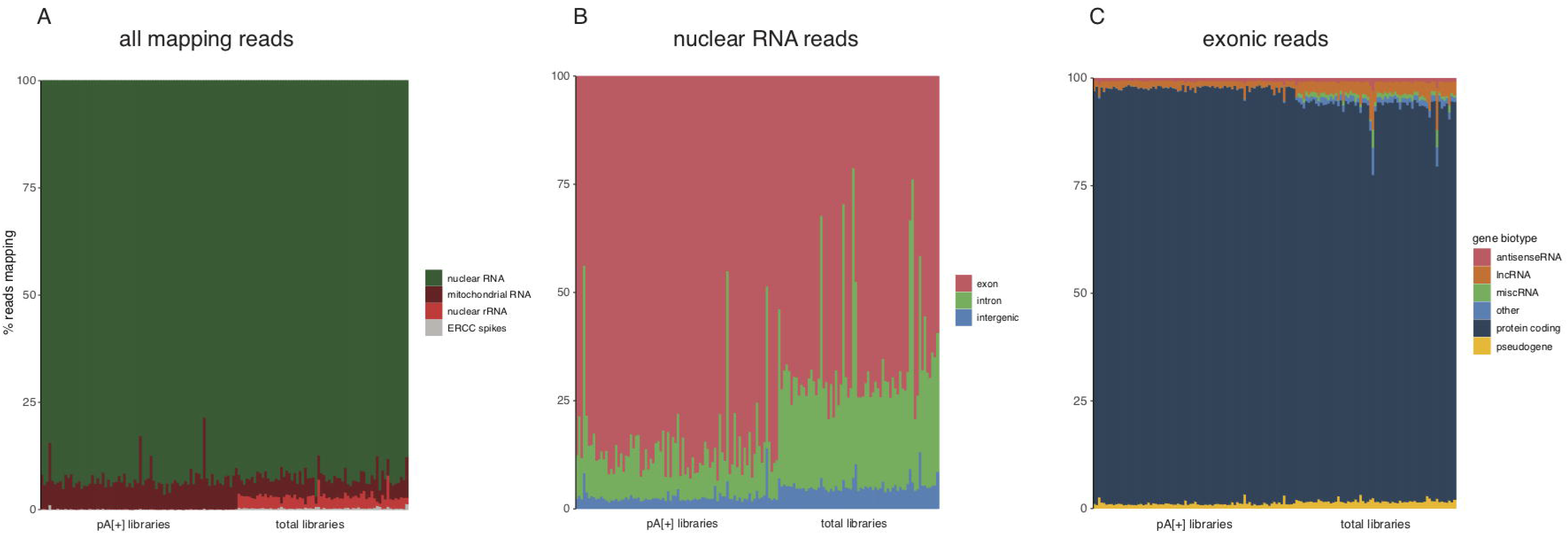
Read distribution differs between polyA[+] and total RNA libraries. A) Percentage of reads derived from nuclear RNA, mitochondrial RNA and ribosomal RNA per cell quantified with STAR. B) Percentage of nuclear reads derived from exonic, intronic and intergenic regions per cell quantified with STAR. C) Percentage of exonic reads attributed to the different biotypes per cell quantified with Kallisto.

### SMARTer single cell total RNA sequencing reveals a unique set of genes

More reads maps to long intergenic RNAs (lincRNAs) using the single cell total RNA-seq protocol (2.64 % [2.523, 2.756; 95 % CI]) compared to polyA[+] RNA sequencing (1.67 % [1.489, 1.849; 95 % CI]). In addition, the single cell total RNA-seq protocol detects an equal or higher number of genes (subsampled to 1 million reads/cell and detected by more than 10 reads) covering the different biotypes, including lincRNAs (144 [139, 148; 95 % CI]), protein coding (5124 [4874, 5372; 95 % CI]) genes, and pseudogenes (132 [127, 137; 95 % CI]) (Figure 2B, 3). Of note, antisense genes are the only biotype for which the total RNA protocol detects fewer genes (62 [59–64; 95 % CI]), likely because of the unstranded nature of the polyA[+] RNA libraries, which results in erroneous quantification of sense/antisense overlapping genes (Supplementary Figure 8). As expected, increasing the number of reads (up to 4 or 8 million) in the total RNA library protocol results in the detection of a higher number of genes. We observed no saturation when generating 8 million reads per cell, suggesting that deeper sequencing could yield even more detected genes (Figure 3). The overlap between protein coding genes detected in the polyA[+] and total RNA libraries (subsampled for 1 million reads/cell and mean expression of at least 1 read over all cells) (Figure 4A) is high. Genes detected in only one of the library types are generally lower abundant compared to genes detected with both methods (Figure 4B). In contrast to protein coding genes, the overlap for lincRNAs between the methods is much smaller (Figure 4C). Importantly, a significant fraction of the total RNA-seq specific lncRNAs display a high expression, thus possibly representing functionally important RNAs (Figure 4D). LincRNA RMRP is one of the most abundant lincRNAs that is solely detected by our novel single cell total RNA-seq workflow. This gene is known to be 3’ nonadenylated and is the first known RNA encoded by a single-copy nuclear gene imported into mitochondria (30, 31). As only a subset of the lincRNAs and antisense genes are currently annotated in Ensembl, we also quantified our libraries with the LNCipedia transcriptome (the most comprehensive human resource of both antisense and lincRNA genes, further referred to as lncRNAs). While the number of detected lncRNAs is slightly lower in the total RNA-seq libraries if an equal number of reads (1 million) is used, each library type contains a certain proportion of unique lncRNAs (Supplementary Figure 9). LNCipedia is likely biased towards medium-to-high abundant polyadenylated lncRNAs.

**Figure 3.**
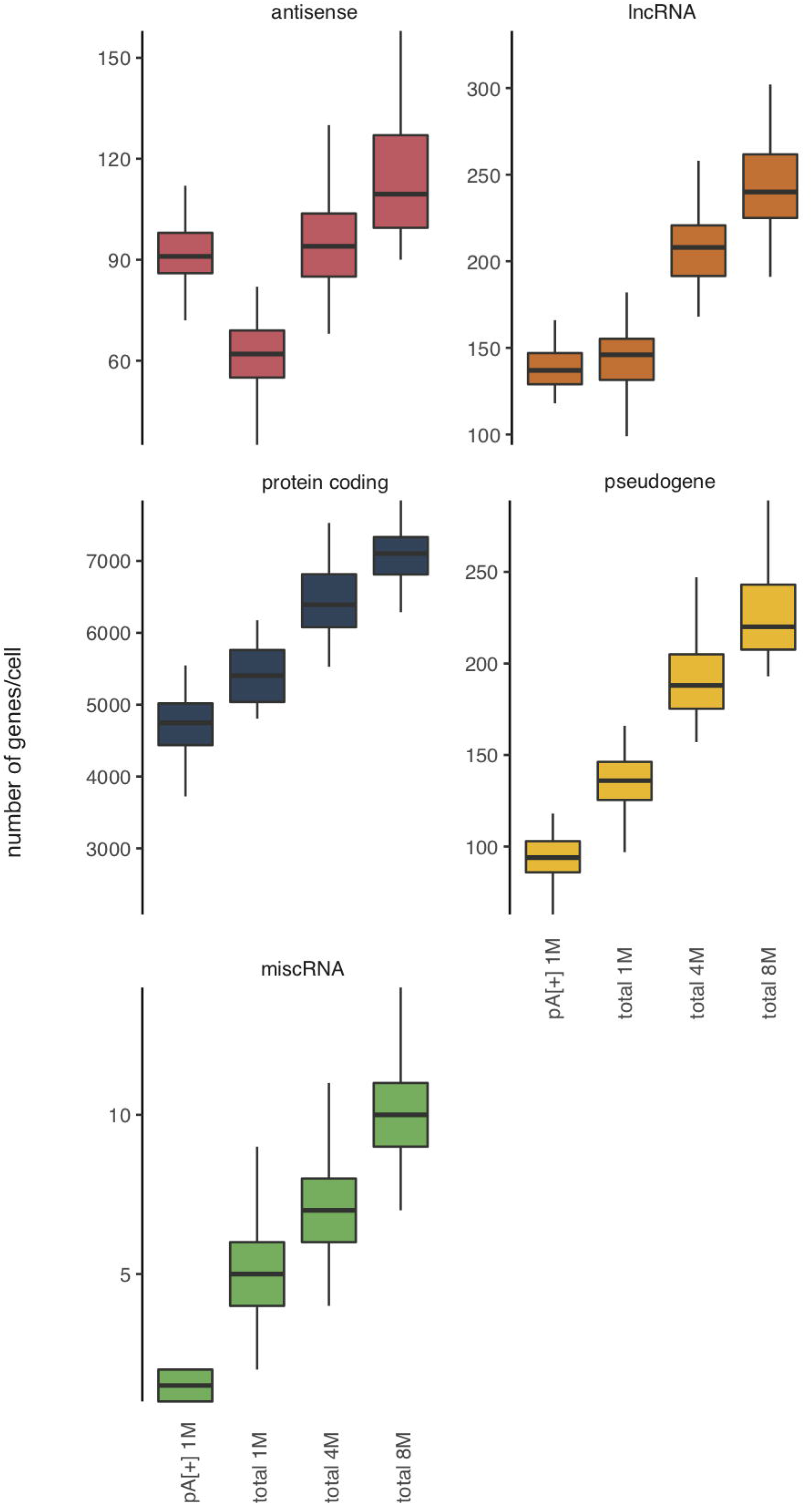
Total RNA libraries detect more genes per biotype. All genes in Ensembl v.91 were quantified on subsampled data (1, 4 or 8 million reads per cell). Only genes with 10 or more reads were included.

**Figure 4.**
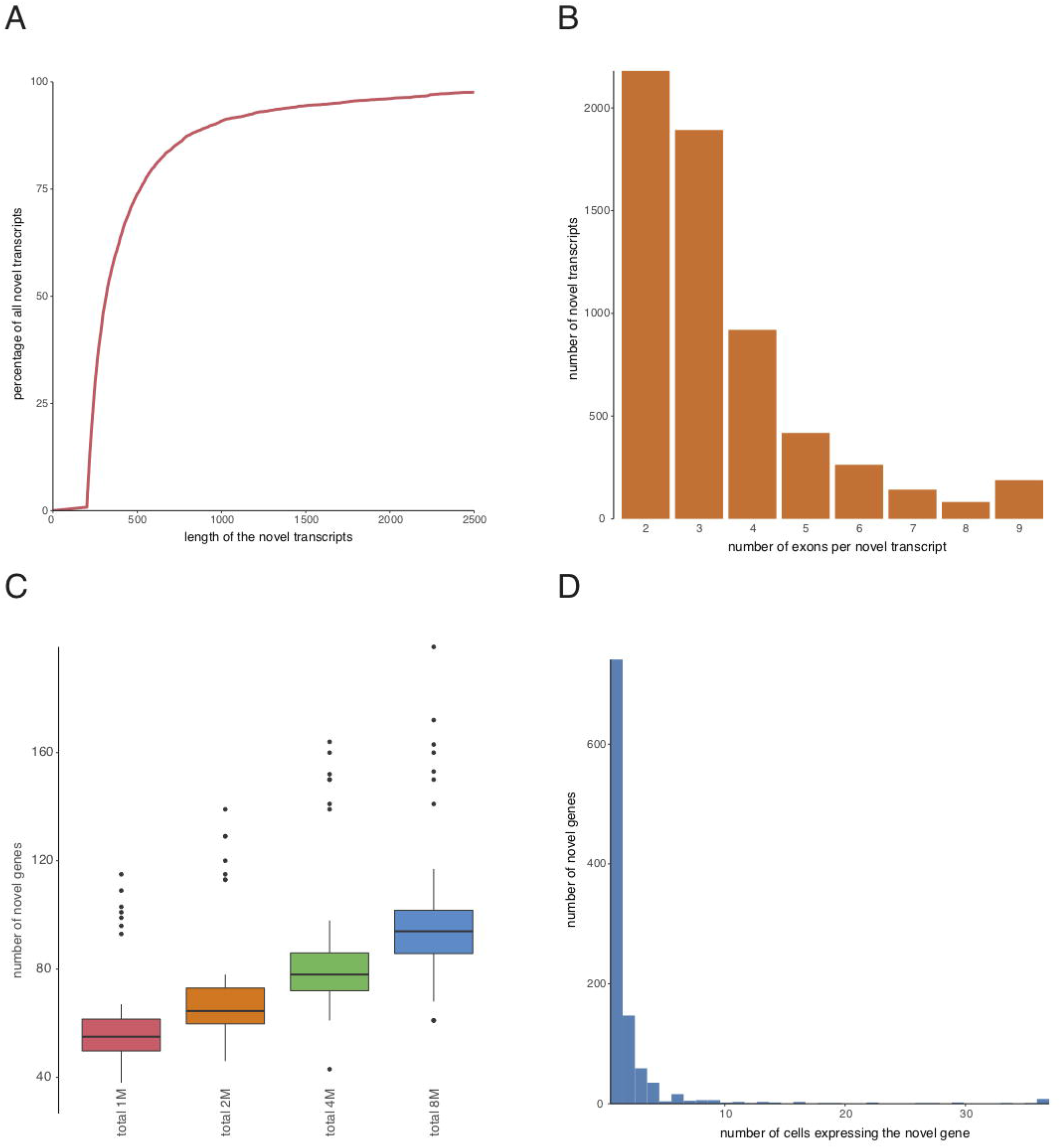
While most protein coding genes are commonly detected, lincRNAs appear more method specific. (A) Overlap between protein coding genes detected in polyA[+] (1 million reads) and total RNA (1 million reads) libraries. (B) Expression counts for protein coding genes detected in only polyA[+] libraries (red), only total RNA libraries (green) or both (gray). (C) Overlap between lncRNAs detected in polyA[+] (1 million reads) and total RNA (1 million reads) libraries. (D) Expression counts for lncRNAs detected in only polyA[+] libraries (red), only total RNA libraries (green) or both (gray).

### SMARTer single cell total RNA sequencing detects circular RNAs and novel genes

In addition to linear RNA biotypes, we tested whether the single cell total RNA-seq protocol is able to quantify circRNAs as this class of non-coding RNAs lacks a polyA-tail and in principle can only be detected using unbiased total RNA-seq. With a requirement of at least two unique back-spliced junction reads, 537 circRNAs were identified derived from 460 host genes (Supplementary Table 2). The majority of the circRNAs were found in fewer than 3 out of 64 cells, with only 14 circRNAs detected in at least 4 out of 64 cells. Gene Ontology analysis for molecular functions and biological processes was performed on the circRNA host genes from both treated and untreated cells. A significant enrichment of TP53 binding, TP53 pathway, cell cycle, and chromosome organization suggests that the identified circRNAs may play a role in these biological functions.

In the single cell total RNA libraries, the fraction of intergenic reads (relative to existing Ensembl and LNCipedia annotation) is high, suggesting that these reads originate from novel unannotated transcripts. To validate this hypothesis, we generated genome and transcriptome guided transcriptome assembly of the paired-end single cell total RNA-seq data resulting in 5360 novel, multi-exonic genes. The novel transcripts have a median length of 317 nucleotides (Figure 5A) and consist on average of more than 3 exons (Figure 5B). Quantification of this novel transcriptome using the single-end data subsampled at 1 million reads per cell resulted in a median number of 59 novel genes per cell [55 – 63; 95 % CI] (Figure 5C). Of note, most novel genes are expressed in only one cell (Figure 5D).

**Figure 5.**
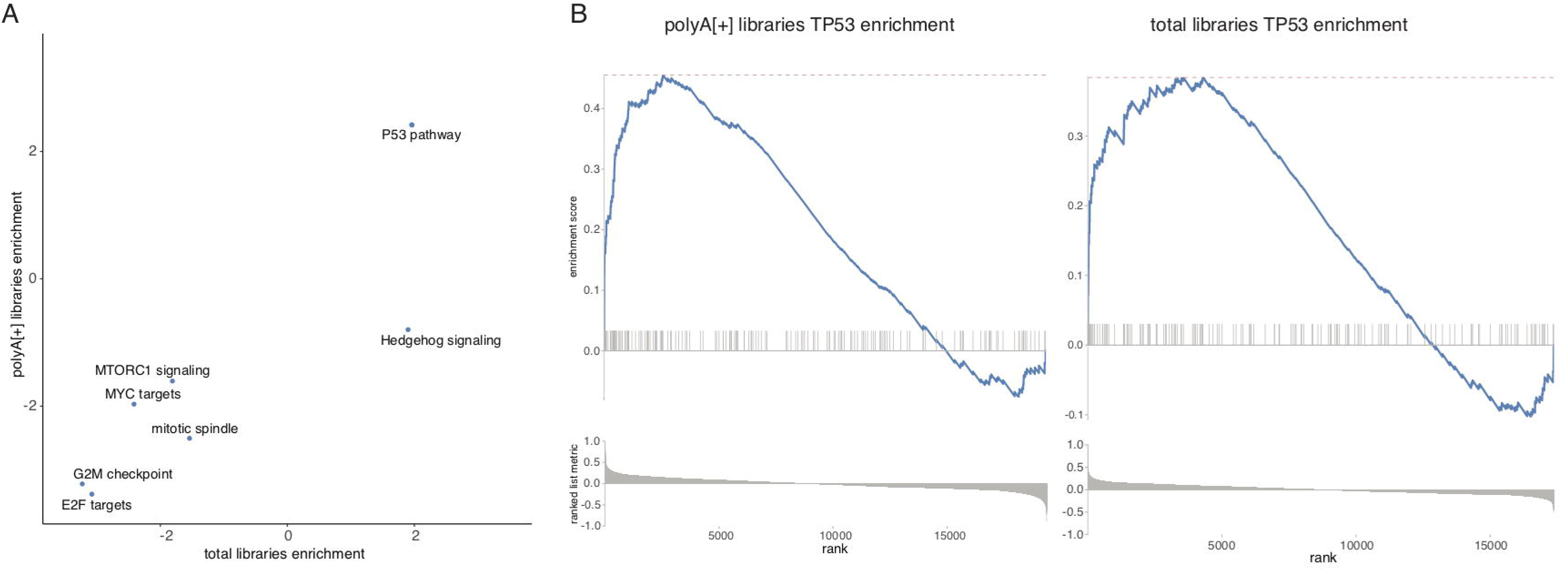
Total RNA libraries enable assembly of single cell transcriptomes. A) Transcripts were filtered at a length of 200 nt. The remaining transcripts have a mean length of 537 nt. B) Transcripts were required to have at least two exons. The remaining transcripts are on average 3.4 exons long. C) All novel genes were quantified on subsampled data (1, 4 or 8 million single-end reads per cell). Genes with at least 1 count were retained. D) While some novel genes are expressed in all cells, most novel genes are detected in only 1 cell.

### SMARTer single cell total RNA profiles reflect the biological signal

To assess whether the single cell total RNA-seq protocol is also able to reveal known biological signal, we performed differential expression analysis using DESeq2 combined with the Zinger method coping with zero inflated data. Based on the ranking obtained by the DESeq2 test statistic, gene set enrichment analysis using the hallmark gene sets was performed. Firstly, the same gene sets are significantly enriched in both library preparation protocols (Figure 6A); secondly, TP53 target genes are – as expected – the most significantly enriched gene set (Figure 6B), confirming that the biological signal is recapitulated through single cell total RNA-seq analyses.

**Figure 6.**
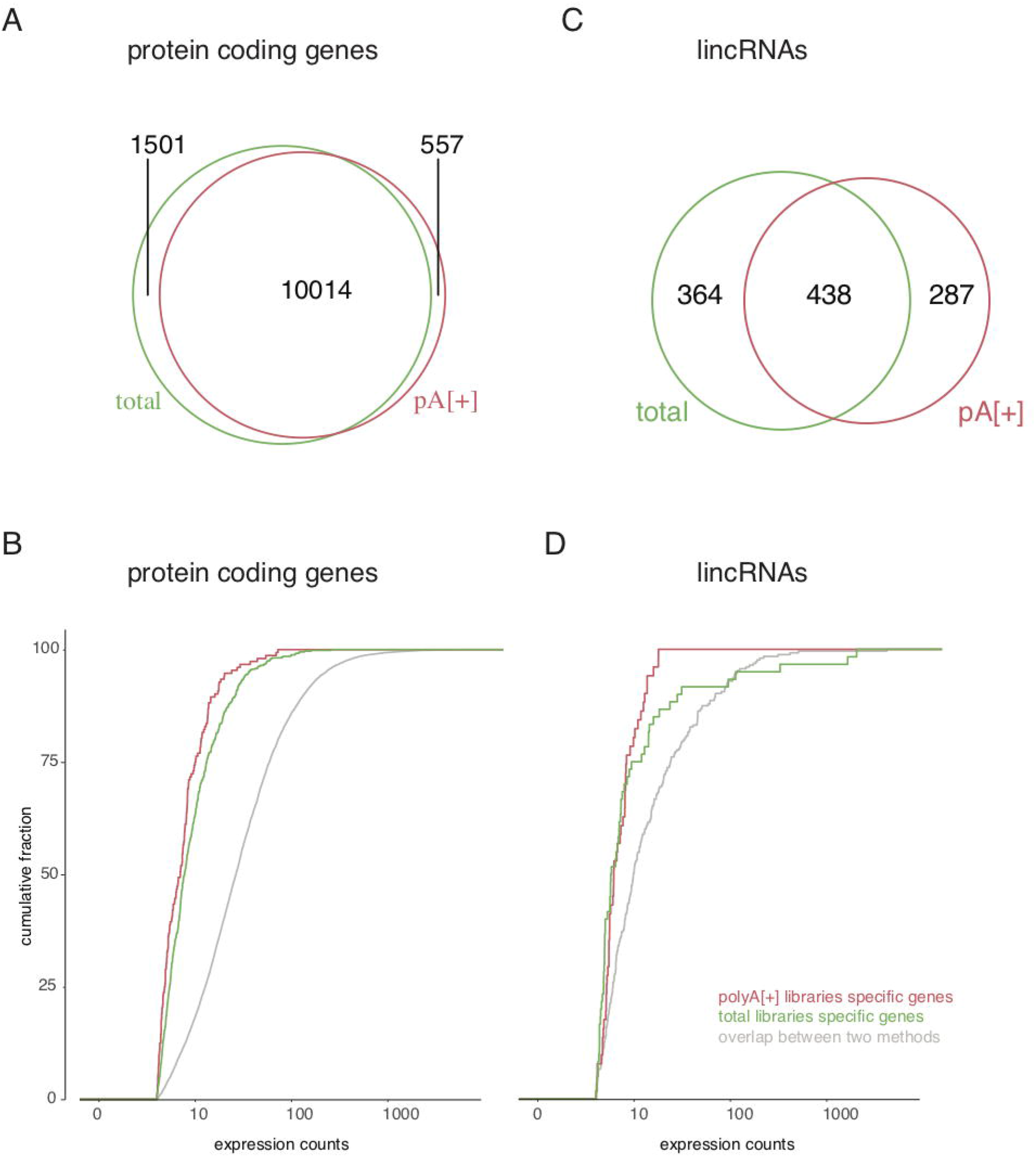
Pathway analysis for polyA[+] RNA and total RNA libraries is similar. A) Gene set enrichment analysis for all hallmark pathways resulted in the same significant (p_adj_ < 0.05) pathway predictions. B) The TP53 pathway is, as expected, enriched in both library prep methods.

## Discussion

In this study, we developed a single cell total RNA-seq method to sequence full transcripts from single cells in an essentially unbiased manner. To demonstrate the performance of the method, we applied single cell total RNA-seq on three different cancer cell lines undergoing a specific perturbation. In parallel, we also performed single cell polyA[+] RNA-seq on the same cells using the well-established SMART-seq method (8). As in any genomics study, the experimental set-up may suffer from confounding factors, such as variations in cell cycle states of the cells and batch effects of single cell capture and sequencing, masking real biological differences. In two of the three experiments, we carefully controlled all these experimental biases. The cell cycle bias was minimized by cell cycle synchronization using serum starvation. We also avoided potential cell selection bias by capturing differentially labeled treated and untreated cells on the same chip (28, 29). Finally, sequencing bias was minimized by sequencing both polyA[+] and total RNA libraries on the same Illumina flow cells.

The single cell total RNA-seq method has some distinctive advantages compared to other methods. First, in any total RNA-seq library, depletion of rRNA is essential as this makes up the bulk of the total RNA mass. Depletion of rRNA from single cells prior to cDNA synthesis is technically impossible. Here, we used ribosomal cDNA specific removal probes, resulting in less than 3 % of ribosomal reads per single cell library. This highly efficient rRNA depletion step is a major improvement compared to RAMDA-seq, where 35 % of the reads map to rRNA (13). Second, as expected, our single cell total RNA libraries contains substantially more intronic reads compared to polyA[+] RNA libraries (32, 33). Such intronic reads can be used to detect changes in nascent transcription, whereby the difference in exonic and intronic reads provides insights in post-transcriptional regulation (34). As such, we believe that our method may be particularly well suited for “RNA velocity analysis” of single cells (35). Third, the single cell total RNA-seq workflow presented in this paper detects relatively more protein coding genes, pseudogenes, lincRNAs and miscellaneous RNA (miscRNA) compared to single cell polyA[+] RNA libraries, when corrected for equal sequencing depth. While the number of detected genes increases with sequencing depth, there seems to be no plateau yet at 8 million reads, suggesting that further increasing the sequencing depth, could enable low abundant gene detection. Fourth, given the stranded nature of the single cell total RNA sequencing data, quantification of antisense genes is accurate, which is not possible when using unstranded data. Fifth, our method also detects non-polyadenylated RNA molecules, such as histone genes, lncRNAs and circRNAs. In the NGP dataset, 537 circRNAs were detected using reads with evidence for back splicing. In order to detect more circRNAs in an individual cell, a higher sequencing depth is required or libraries should be enriched for circRNAs by selectively removing linear RNA by exonuclease treatment prior to library prep and sequencing (11, 12). Sixth, the data enables reference guided transcriptome assembly, resulting in 5360 novel genes. Finally, differential gene expression analysis and gene set enrichment of NGP cells treated with nutlin-3 confirmed activation of the TP53 pathway at the transcriptional level.

One limitation of the implementation of the single cell total RNA library preparation method on the C1 instrument is the relatively low throughput, as maximally 96 cells are simultaneously captured. In contrast, current droplet based single cell methods capture thousands of individual cells, but these systems are limited to 3’ end sequencing of polyadenylated RNA, preventing quantification of splice variants and non-polyadenylated transcripts. Higher cell numbers as input for the total RNA library prep method can in principle be achieved by using FACS technology. Our advice is to use the single cell total RNA-seq method rather than polyA[+] methods if it is desired to study non-polyadenylated RNA molecules such as lncRNAs or circRNAs, if stranded-specific data is a must and if full transcript sequencing is priority (e.g. analysis of alternative splicing, RNA editing or somatic mutations).

## Availability

The SMARTer single cell total RNA sequencing script is deposited in Script Hub (Fluidigm).

## Accession Numbers

The fastq files and processed data is available through GEO (GSE119984).

To review GEO accession GSE119984:

Go to https://www.ncbi.nlm.nih.gov/geo/query/acc.cgi?acc=GSE119984 Enter token orepemiibzedruh into the box.

## Supplementary Data

Supplementary Data are available at NAR online.

## Author Contributions

K.V. ported the bulk library prep method to a single cell library prep method on the C1, with help from N.B and K.J.L. C.E. analyzed the data. K.V., C.E. and J.V. wrote the manuscript. K.V. and N.Y. performed the experiments on the C1 system. D.R. assisted with the serum starvation experiment and the experiments on the SHSY5Y-MYCN-TR cell line. J.A., M.T.V. and J.K. performed circRNA analysis. F.S., P.M. and J.V. supervised the project. All authors approved the final version of the manuscript.

